# DYRK1A up-regulation specifically impairs a presynaptic form of long-term potentiation

**DOI:** 10.1101/645200

**Authors:** Aude-Marie Lepagnol-Bestel, Simon Haziza, Julia Viard, Paul A. Salin, Valérie Crépel, Arnaud Duchon, Yann Herault, Michel Simonneau

**Author notes:** Correspondence: Simonneau M.

## Abstract

Chromosome 21 DYRK1A kinase has long been associated with a variety of psychiatric diseases including Down Syndrome. We previously showed that Dyrk1A interacts with SWI/SNF (SWItch/Sucrose Non-Fermentable) nucleosome remodeling complex inducing expression changes of genes encoding key neuronal proteins. However, the functional impact of this kinase at the synapse level remains unclear. We studied a mouse model that incorporated the YAC 152F7 (570 kb) encoding six chromosome 21 genes including DYRK1A. We found that DYRK1A Interacts with the key chromatin remodelers EP300 and CREBBP. Moreover, we observed changes in the transcriptional levels of genes encoding presynaptic proteins involved in glutamate vesicle exocytosis, namely Rims1, Munc13-1, Syn2, Rab3A. This result prompted us to investigate the two main forms of long-term potentiation (LTP) required for learning and memory: the (N-methyl d-aspartate) receptor-dependent postsynaptic form versus the glutamate release-dependent presynaptic form. Interestingly, extracellular electrophysiological recordings in hippocampal slices of the YAC mouse line revealed that only the presynaptic forms of plasticity were impacted, leaving the post-synaptic form of plasticity intact. T o refine our findings, we used a mouse BAC 189N3 (152 kb) line that only triplicate the gene Dyrk1A. Again, we found that this presynaptic form of LTP is also impaired in this mouse line. This result demonstrates that abnormal up-regulation of Dyrk1A alone is sufficient to inhibit specifically the presynaptic forms of LTP. Altogether, our results suggest that impairment of DYRK1A gene dosage may impact memory precision, and therefore reinforce our mechanistic understanding of the cognitive impairment detected in this mouse model.

## INTRODUCTION

Down syndrome (DS), a human genetic disorder, is the most common form of intellectual disability (ID), affecting 1/1,000 births. DS is caused by the presence of a third copy of up to ~234 genes from *Homo sapiens* autosome 21 (Hsa21) (1). Despite a broad spectrum of clinical symptoms, virtually all people affected by DS develop Alzheimer’s disease pathology by 40 years of age, with intellectual deficits that impair learning and memory (1–4). However, a detailed understanding of the precise contribution of each Hsa21 gene to the cognitive impairment found in DS patients is still lacking.

Chromosome 21 Dyrk1A (dual-specificity tyrosine phosphorylated and regulated kinase 1A) gene is a major player in DS whom overexpression impacts the synaptic plasticity within the hippocampus and the prefrontal cortex (5–7). Converging evidence suggests that DYRK1A might be critical for learning and memory processes (8, 9). However, a clear understanding of the regulatory pathways impaired by a trisomy of DYRK1A is still lacking.

Here, we used two DS mouse models. The first model is a human YAC mouse model (152F7 line) that displays a ~570 kb human genomic region surrounding the *DYRK1A* gene and including five other genes (10). The second is a Dyrk1A BAC model (189N3 line) that gives a triplication of ~152 kb mouse Dyrk1a locus (6). By performing exome sequencing and mass spectroscopy in these mouse models, we recently uncovered two deregulated repertoires associated with chromatin and synaptic pathways (11). Interestingly, several studies reported that these lines display behavioral defects in episodic-like forms of memory (10, 12) Indeed, both 152F7 and 189N3 mouse models showed impairment in long-term spatial memory tested by Morris Water Maze (10, 12, 13) or Novel Object Recognition task (14).These results indicate that both the “where” and the “what” pathways of episodic memory (15) are impacted in these models. These two pathways involve respectively the Lateral Entorhinal Cortex (LEC) that encodes experience from a first-person (egocentric) perspective, and the Medial Entorhinal Cortex (MEC) to the world-centered (allocentric) coding of place cells, grid cells, and head direction cells (16). Both LEC and MEC pathways share the same Dentate Gyrus-CA3 (Mossy Fiber-CA3) and CA3-CA1 pathways (15). In spite of these defects in both “where” and “what” hippocampal pathways, no study of long-term potentiation (LTP), a prominent cellular model thought to underlie long-term memory, have been reported for both 152F7 and 189N3 mouse lines (6, 10).

In this paper, our aim was to evidence a relation between these cognitive defects and a functional synaptic defect in these 152F7 and 189N3 mouse lines. Here, we first show here that DYRK1A directly interacts with two chromatin remodelers in neuron nuclei: EP300 and CREBBP. Moreover, using in situ hybridization, we found that deregulation of DYRK1A dosage indirectly induced changes in expression of genes encoding pre-synaptic proteins. Altogether, these results prompted us to investigate whether the synaptic plasticity is impaired in the Dentate Gyrus Mossy Fiber-CA3 pathway that is known to involve a presynaptic form of LTP linked to presynaptic proteins (17, 18). Therefore, using extracellular field recording in hippocampal slices, we found that the NMDA-independent long-term potentiation of the Dentate Gyrus-CA3 synapses is specifically impaired in these two mouse lines. These results reveal a previously unappreciated impact of Hsa21 DYRK1A on synaptic plasticity with possible consequences for a variety of neuropsychiatric diseases as diverse as Intellectual Disability, Late-Onset-Alzheimer Disease and Autism Spectrum Disorders.

## RESULTS

### Hsa21 DYRK1A interacts with the chromatin remodelers CREBBP and EP300

The 152F7 mouse model incorporates 570 kb fragment of Hsa21 with six protein-coding genes including DYRK1A (**Supplementary Figure 1**). One of the six genes, *DSCR9* is a primate-specific gene (19). We recently evidenced (11) that DSCR9 directly interacts with clusterin, a genetic risk factor of Late-Onset Alzheimer disease (20). The syntenic region in mouse chromosome 16 genome is more condensed with ~350 kb (**Supplementary Figure 2**) instead of ~570 kb in Hsa21. The organization of genes is similar between human and mouse genome with the sequence 3’ to 5’: Ripply3, Pigp, TTC3, Vps26c and Dyrk1A.

Here, we first analyzed protein-protein interactions in order to unravel possible molecular changes involved in cognition. We recently uncovered DS-related protein-protein interaction (PPI) map using a high-throughput, domain-based yeast two-hybrid (Y2H) technology against a human brain library (11). We attained more than 1200 PPI, among which we identified direct interactions between DYRK1A and EP300 or CREBBP (11). This result is consistent with previous findings using mass spectrometry methodologies (21). EP300 and CREBPP are two members of the p300-CBP coactivator family containing a histone acetyltransferase (HAT) domain involved in chromatin remodeling (22). Mutations in these genes have been shown to cause Rubinstein-Taybi Syndrome (23). To further validate these two interactions, we performed immunoprecipitation (IP) on HEK293 cells using anti-EP300 and anti-CREBBP antibodies and successfully identified DYRK1A (**Figure 1A-B**). We next validated direct nuclear interactions of Dyrk1A with EP300 and CREBBP in cortical neurons *in vitro* using a proximity ligation assay (PLA) (**Figure 1C** and **Supplementary Figure 3**).

**Figure 1.**
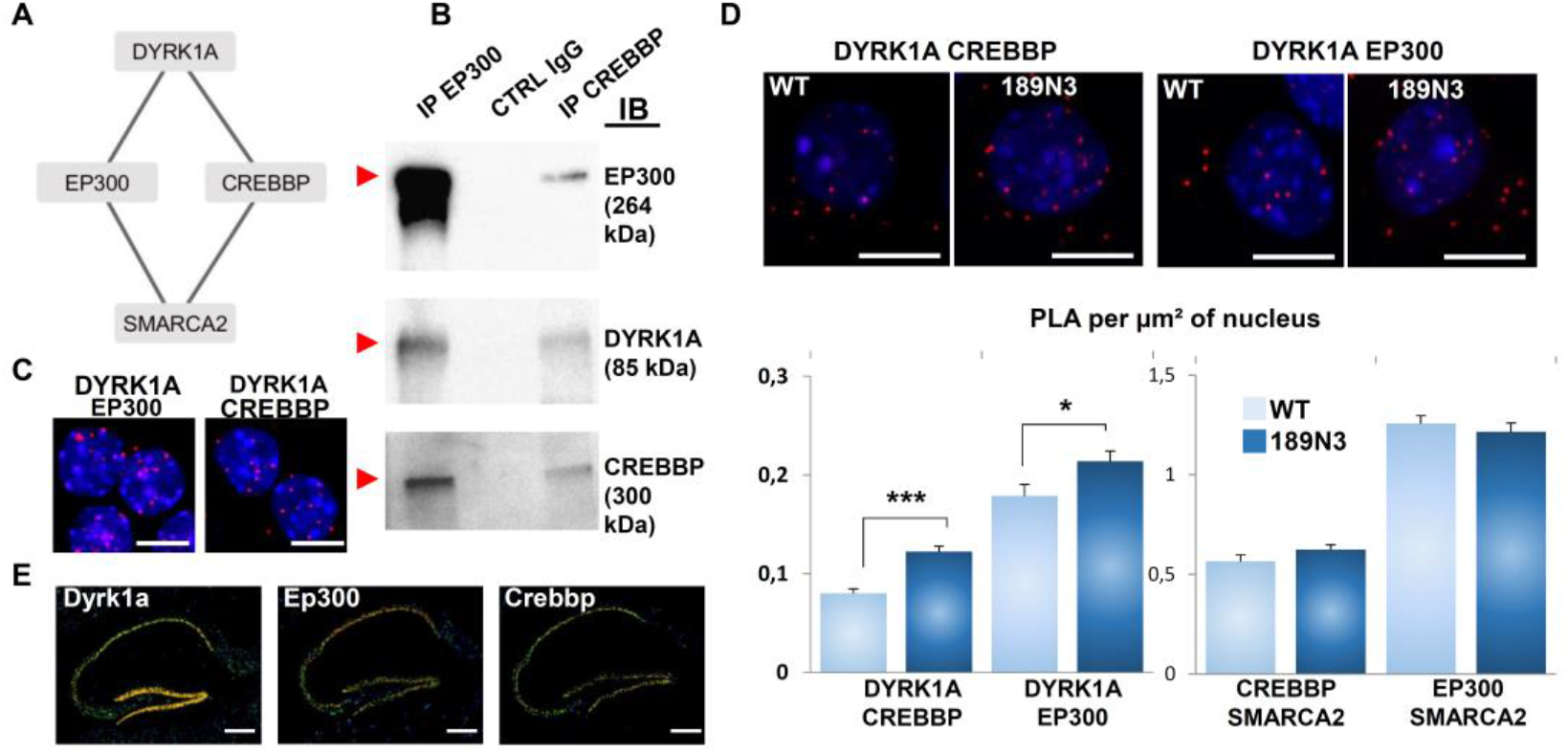
Interaction of HSA21 DYRK1A with chromatin remodelers. A. Schematic representation of DYRK1A interaction with EP300 and CREBBP. B. HEK293 cells were immunoprecipitated (IP) using anti-EP300 and anti-CREBBP antibodies and using anti-IgG antiboby as a negative control. The input and precipitated fractions were analyzed by western blot using anti-Ep300, anti-Dyrk1a and anti-Crebbp antibodies. The arrows indicate proteic bands at the expected size. Note that no cross-reaction was found with the IgGs. C-D. *In situ* proximity ligation assays (PLA) on primary cortical neurons fixed at DIC7 (red fluorescence) using anti-Dyrk1a and anti-Ep300 or anti-Crebbp, anti-Smarca2 and anti-Ep300 or anti-Crebbp antibodies. Nuclear bodies were labelled using Topro3 staining (blue fluorescence). Mean interaction point numbers were calculated in nuclear body of 45 to 89 cortical neurons at DIC7 (from 3 to 5 different embryos per genotype). PLA using anti-Ep300 and anti-Fibrillarin antibodies were performed as a negative control and no difference was shown between transgenic 189N3 and WT cortical neurons. Scale bars=10μm. * p < 0.05; *** p < 0.0005. E. False-color image of ISH from Allen Brain Atlas showing Dyrk1a, Ep300 and Crebbp transcript expression in adult mouse hippocampus. Scale bar = 100μm.

Quantification of these interactions in 189N3 transgenic neurons compared to wild-type neurons indicate a statistically significant increase in DYRK1A-EP300 and DYRK1A-CREBBP interactions (**Figure 1D**). In contrast, we found no changes in the number of EP300-SMARCA2 and CREBBP-SMARCA2 interactions between 189N3 transgenic neurons compared to wild-type neurons (**Figure 1D** and **Supplementary Figure 3**). These results suggest that this increase in the number of DYRK1A-EP300 and DYRK1A-CREBBP in 189N3 can induce changes in transcriptome of neurons. Quantitative *In Situ* Hybridization (ISH) from Allen Brain Atlas indicates that both *Dyrk1a, Ep300* and *Crebbp* are highly expressed in adult mouse hippocampus (**Figure 1E**). Altogether, these results identify chromatin regulators as direct interactors of DYRK1A.

### Dyrk1A up-regulation affects the transcription of genes encoding presynaptic proteins

As *EP300* and *CREBBP* encode HATs and impact chromatin remodeling, we hypothesized that changes in the level of DYRK1A would affect its interactions with these HAT partners and would consequently modify the transcription of genes involved in synaptic function (24–27). We tested whether genes encoding presynaptic proteins were deregulated in 152F7 hippocampal sub-regions. We first quantified Rims1 transcriptional changes using quantitative radioactive ISH on hippocampal sections of 152F7 and wild type (WT) mice. We detected a significant decrease in Rims1 transcript in both DG, CA3 and CA1 (**Supplementary Figure 4**). Both Rim1, Rab3a, Rims1 and Munc13-1 are involved in the presynaptic release of glutamate (28) (**Figure 2A**). Synapsin I and synapsin II are widely expressed synaptic vesicle phosphoproteins that have been proposed to play an important role in synaptic transmission and short-term synaptic plasticity (29, 30). We first used Western blotting of Rim1, Rab3a, Munc13-1 and Synapsin 2A proteins from hippocampus extracts of 152F7 and WT mice. We found a decrease in Synapsin 2A protein for 152F7 as compared to WT mice (**Figure 2B**). We next used RT-Q-PCR following laser-assisted micro-dissection of hippocampal sub-regions (**Figure 2C**) to quantify possible changes of these transcripts between 152F7 and control mice. We found that the protein levels of Rims1, Syn2, Munc13-1 and Rab3a decreased in 152F7 dentate gyrus compared to wild type (**Figure 2D**). We compared transcript levels for dorsal DG, CA3 and CA1 from our data obtained by Q-RT-PCR from laser-assisted microdissection with Hipposeq data obtained and from single-cell transcriptomics (31). Quantitative changes were fully coherent for Rims1 and partially for Syn2, not coherent for Rab3a and Munc13-1, suggesting 3-D heterogeneity (**Supplementary Figure 5**).

**Figure 2.**
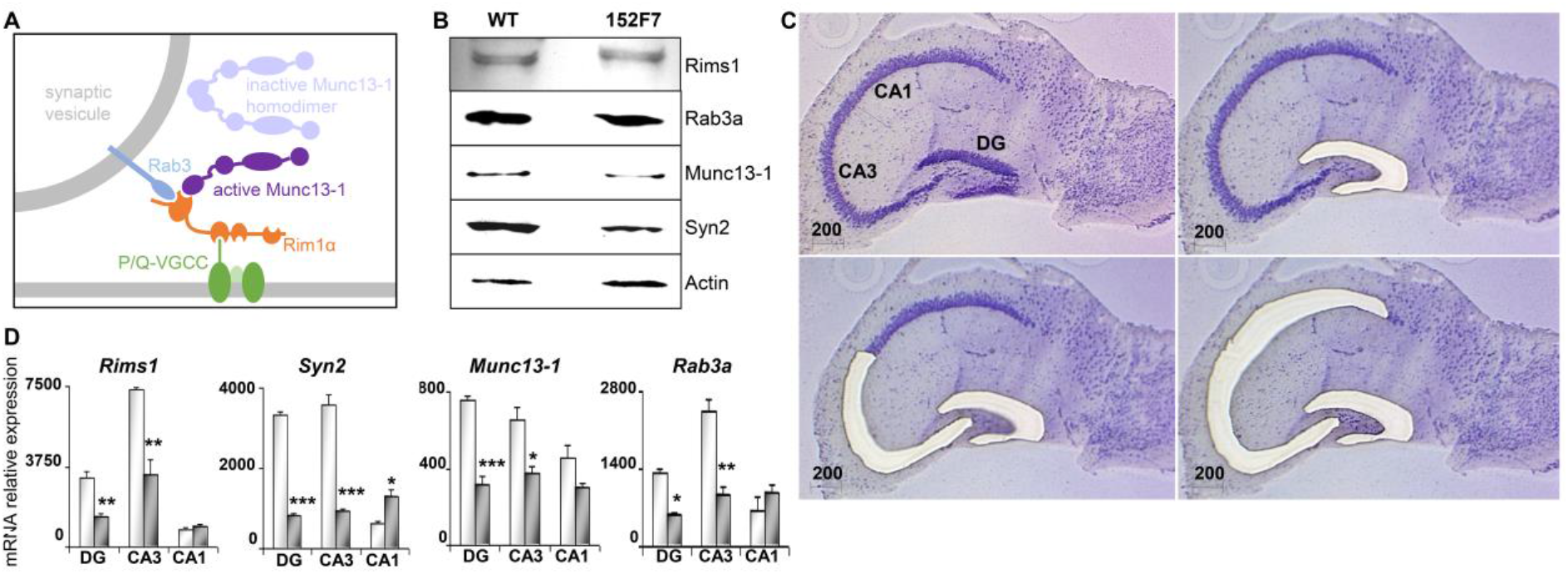
Presynaptic protein expression in the adult 152F7 mouse hippocampus as compared to control mice. A. Schematic representation of molecules involved in glutamate release from presynaptic vesicles. B. Western blotting of Rim1, Rab3a, Munc13-1 and Synapsin 2A proteins from hippocampus extracts of wild-type and 152F7 transgenic mice. The amounts of protein were normalised to actin levels. Four to five hippocampi were pooled. There was a decrease in the amount of Synapsin 2A in the 152F7 transgenic mice (152F7) as compared to the wild type (WT). C & D. Laser-assisted microdissection of the three subregions of P21 mouse hippocampus stained with toluidine blue (C). Scale bar = 200 μm. From Laser-assisted microdissection of the three subregions of P21 mouse hippocampus, *Rims1, Syn2, Rab3a* and *Munc13-1* transcripts are down regulated in the DG and CA3 hippocampal subregions of juvenile transgenic 152F7 mice compared to their WT siblings as shown by Q-RT-PCR analysis (D). Scale bar = 1mm. *p<0.01**p<0.001. ***p<0.0001

Altogether, these results show that change in *Dyrk1a* dosage can modify the expression of gene encoding proteins involved in the molecular mechanism of presynaptic vesicle release.

### Dyrk1A up-regulation specifically impairs a presynaptic form of LTP at the DG-CA3 hippocampal synapse

Two main forms of LTP have been characterized in the mammalian brain. One requires activation of postsynaptic NMDA (N-methyl d-aspartate) receptors, whereas the other, as for mossy fiber (MF) LTP, is NMDAR independent and require presynaptic mechanisms. This presynaptic form of LTP at the MF-CA3 synapse have been widely characterized and requires activation of several presynaptic proteins linked to glutamate vesicle exocytosis such as Rims1, Munc13-1, Syn2, Rab3A (17, 28).

Here, we used field excitatory postsynaptic potential (fEPSP) to functionally test the effect of abnormal up-regulation of Dyrk1a in both mouse models 152F7 and 189N. We recorded in the lucidum of CA3 and MF-mediated fEPSP were evoked by weak electrical stimulations in the dentate gyrus. We induced LTP by tetanus at 25Hz during 5s under NMDA blocker D-AP5 (D-2-amino-5-phosphonovalerate) to prevent NMDA-dependent LTP from occurring (**Figure 3 A**). In both models, we found that this form of NMDA independent LTP was absent at the dentate gyrus-CA3 synapse (**Figure 3 B-D**), suggesting that Dyrk1A overexpression alone is sufficient to inhibit mossy fiber LTP. These results show that abnormal *Dyrk1a* dosage can modify the expression of genes encoding proteins involved in the molecular mechanism leading to presynaptic release.

**Figure 3.**
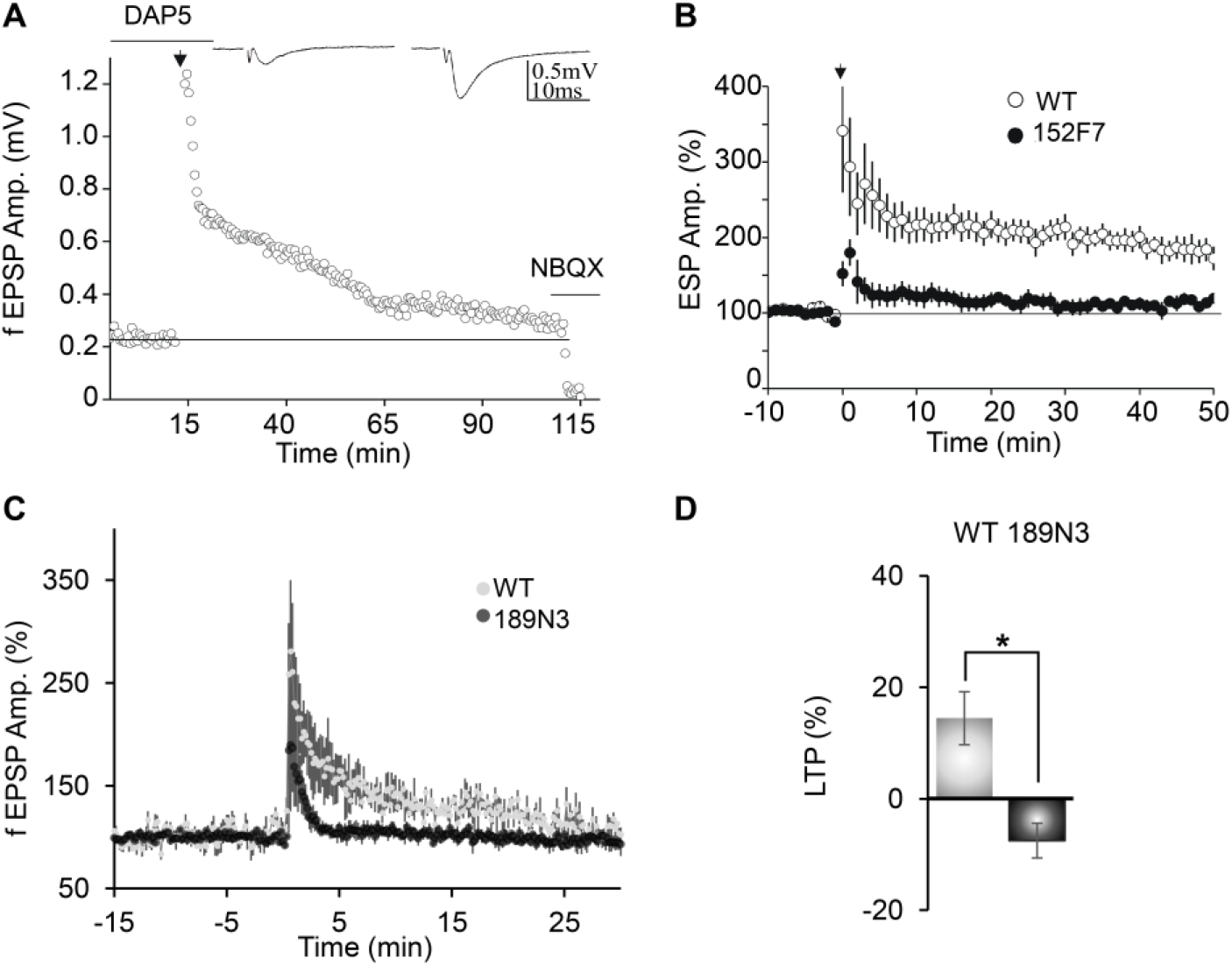
Presynaptic LTP between Dentate Gyrus mossy fibers and CA3 impaired in both adult 152F7 and 189N3 mouse hippocampus as compared to control mice. A. Example of a temporal trace of mossy fiber LTP in a wild type mouse. Mossy fiber LTP was induced by a single tetanus of 25 Hz (for 5 s, black arrow) in the presence of 50 μM of DAP 5. NBQX was applied at the end to obtain information concerning the fiber volley. Inset: example traces of a baseline fEPSP (left) and fEPSP after LTP (right). B. Summary of mossy fiber LTP in transgenic 152F7 compared to WT mice, showing LTP induction defect in the DS mouse model. Data are means+/ SEM and calculated from 4 different mice with 3 slices per mouse for both genotypes. C. D. Summary of mossy fiber LTP in transgenic 189N3 compared to WT mice (C). Average fEPSP amplitude 20 min after LTP induction in WT versus 189N3 mice, showing LTP induction defect in the DS mouse model (D). Recording were performed on 4 to 6 mice in each genotypes. Data are means+/ SEM.

## DISCUSSION

The general aim of this study was to evidence a relation between the cognitive defects and synapse plasticity impairments in the 152F7 Down syndrome mouse model (10), in which episodic spatial memory is impaired without noticeable defects in hippocampal synaptic plasticity (10, 13). The 152F7 mouse model expresses a human YAC fragment of 570 kb that includes six distinct genes, among which two transcription regulators, RIPPLY3 and DYRK1A. RIPPLY3 was characterized as a transcriptional co-repressor but its putative function is far from being understood. In contrast, *DYRK1A* encodes *a* kinase with multiple subcellular sites of action including transcriptional regulation. We previously demonstrated that DYRK1A interacts with the REST/NRSF-SWI/SNF chromatin remodeling complex to deregulate gene clusters involved in the neuronal phenotypic traits of Down syndrome (32). Moreover, it was recently shown that the DYRK1A regulates transcription of a subset of genes by associating to their proximal promoter regions and phosphorylating the C-terminal domain of the RNA polymerase II (33). In this study, we found that DYRK1A directly interacts with chromatin remodelers EP300 and CREBPP that are known to impact synaptic plasticity (24).

Recent data suggest that memory precision defects can be generated by deregulation of genes involved in hippocampus Dentate Gyrus-CA3 connections, in particular Dentate Gyrus Mossy Fibers (MF)-CA3 synapse (18, 34–38). A During contextual fear memory consolidation, the strength of mossy fiber connections is selectively increased between dentate gyrus engram cells and CA3 engram cells (i.e. mossy fiber LTP-like process, Ryan et al, 2015). This interplay between the dentate gyrus and CA3 area is required for long-term memory process that requires pattern separation, a process that inhibits memory generalization. Increased time-dependent loss of memory precision is characterized by an overgeneralization of fear that is a feature of post-traumatic stress disorder (PTSD) (39). Memory imprecision also characterizes mild cognitive impairment (MCI) that precedes Late-Onset Alzheimer Disease (40, 41). The connectivity between Dentate gyrus cells (DGC) and CA3 pyramidal neurons via DGC MF in the hippocampus is central for memory precision (32, 34). Hippocampal MF are the presynaptic component of MF-CA3 synapses that display both structural and functional plasticity in relation to the precision of hippocampus dependent memories. The earliest indication that large presynaptic boutons called large mossy fiber terminals (LMTs) have distinct physiological properties originated from the finding that the induction of LTP at LMTs was independent of NMDA receptor activation (42). Since MF-LTP was induced independently of activity on the postsynaptic site, it was suggested that this LTP was solely triggered by the presynaptic terminal itself (43). The presynaptic vesicle protein Rab3a and the active zone protein Rims1 and Munc13-1 have been shown to be required molecular components of the presynaptic mechanism underlying the maintenance and expression of MF-LTP in the hippocampus (44–47)

From these results, we examined if *DYRK1A* gene dosage can modify the regulation of genes that encode proteins involved in presynapse vesicle exocytosis as knockout of the related genes disrupt MF-CA3 non-NMDA LTP. Using quantitative ISH, we were able to find a statistically significant decrease of *Rims1* in 152F7 Dentate Gyrus. We identified significant decrease in transcripts level involved in presynapse vesicle exocytosis such as Rims1, Rab3a and Munc13-1 whose knockout induces impairment in MF-CA3 LTP. Other transcripts such as Syn2 involved in presynapse functioning (Spillane et al., 1995) were also impacted. Impairment of MF-CA3 LTP has been reported only for knockout of Rims1, Rab3 and Munc13-1 with no effect for haploinsufficiency (44–47). Here, we propose that the combined haploinsufficiency for Rims1, Rab3 and Munc13-1 can have a functional consequence similar to knockout of one of these genes, leading to a phenocopy of knockout.

Furthermore, we were able to demonstrate using the 189N3 mouse model overexpressing only Dyrk1A that MF-CA3 LTP defect can be directly linked to Dyrk1A overexpression. Implication of DYRK1A in LTP defects that is linked to memory precision can be of clinical importance for Down Syndrome but also for Autism Spectrum Disorders as DYRK1A is one of the key candidate genes in this disease with a ~20 fold increases in risk (48, 49). Altogether, this work identifies novel molecular and functional novel biomarkers that are involved in the cognitive signature displayed by mouse models of Down Syndrome. Our study thereby provides a novel mechanistic and potentially therapeutic understanding of deregulated signaling downstream of DYRK1A up-regulation.

## ACKNOWLEGDMENTS

This work was partly supported by European FP7-HEALTH AgedBrainSysbio and European JPND TransPathND program (to M.S.).

## MATERIALS AND METHODS

### Animals and genotyping

We used wild-type mice of the OF1 strain for neuronal primary culture, of the C57BL6 strain and 189N3 or Dp(16)1Yey transgenic line for neuronal primary cultures and electrophysiology and FVB strain and the 152F7 transgenic line for electrophysiology (10). Genotypes were determined using genomic DNA extracted from skeletal muscle fragments and the PCR protocol and primers as described previously (32).

### Quantitative *In situ* hybridization (ISH)

For Rim1 *in situ* hybridization, a 1124 bp Rim1 PCR product (covering nucleotides 376 to 1499 of the XM_129709 cDNA sequence) was inserted into a pCRII-TOPO cloning vector) as described in. XhoI-or HindIII-linearized Rim1 inserts were used to generate antisense or sense [α35S]-rUTP (800 Ci/mmol, Amersham) labeled transcripts, using the P1460 riboprobe in vitro transcription systems (Promega), according to the manufacturer’s instructions. Paraffin-embedded coronal 15 μm sections of P21 mouse brains, including hippocampal structures, were selected for ISH. Sections were hybridized with radiolabeled RNA probe (diluted to 105 cpm/μl in 50% formamide hybridization buffer) in 50% formamide at 50°C. Sections were successively washed in 50% formamide, 2 x SSC, 10 mM DTT at 65°C, and then in increasingly stringent SSC washing solution, with a final wash at 0.1 x SSC at 37°C. The regional distribution of radioactivity was analyzed in the hippocampal region of wild-type and transgenic sections using a digital autoradiography imager (Micro Imager from Biospace Lab, Paris, France) and its analysis program (BetaVision, Biospace Lab).

### Protein extraction and Western blot analysis

HEK293 cells or mouse cortex (pool from three adult OF1 mice) were homogenized on ice in Tris-buffered saline (100 mM NaCl, 20 mM Tris-HCl, pH 7.4, 1% NP40, 1 x CIP). The homogenates were centrifuged at 13,000g for 10 min at 4°C and the supernatants were stored at −80 °C. Cell lysate protein concentration was determined using the BCA Protein assay kit (ThermoFisher). For SDS-PAGE, 40μg of protein was diluted in Laemmli 1x (BioRad) with DTT and incubate for 5mn at 95°C. Proteic samples were loaded in each lane of a 4-15% precast polyacrylamide gel (BioRad) and ran in Mini-Protean at 200 V in Tris/Glycine running buffer (BioRad). Following SDS-PAGE, proteins were semi-dry electroblotted onto nitrocellulose membranes using the Trans-Blot Turbo Transfer System (BioRad). Membranes were incubated for 1 h at room temperature in blocking solution (PBS 1x containing 5% non-fat dried milk, 0.05% Tween 20) and then for overnight at 4°C with the primary antibody. Primary antibodies used were as shown in Supplementary Table S6. Membranes were washed in PBS 1x containing 0.05% tween 20 and incubated for 1 h at room temperature with anti-mouse, anti-rabbit or anti-goat HRP-conjugated secondary antibody. Membranes were washed three times in PBS 1x containing 0.05% tween 20. Immune complexes were visualized using the Clarity Western ECL Substrate (BioRad). Chemiluminescence was detected using the ChemiDoc XRS Imaging System (BioRad). Primary antibodies used were as shown in Supplementary Table S3. As secondary antibodies, we used protein A or protein G IgG, HRP-conjugated whole antibody (1/5,000; Abcam ab7460 or ab7456 respectively).

### Laser-assisted microdissection, Total RNA preparation and quantitative real-time PCR (Q-RT-PCR) analysis

Embryonic brain subregions were dissected as shown in Supplementary Figure. The left and right hippocampus was micro dissected from genotyped P21 mouse brains using a laser-assisted capture microscope (Leica ASLMD instrument) with Leica polyethylene naphthalate membrane slides as described in. RNA preparation and Q-RT-PCR are performed as described in. Q-RT-PCR results are expressed in arbitrary unit.

### Electrophysiological analysis on 152F7 juvenile mice

The methods used have been described elsewhere (Hanson et al., 2007). Briefly, for the preparation of hippocampal slices, 21-day-old mice were deeply anaesthetized with Nembutal. Brain slices (300–400 μm) were cut in cold artificial cerebrospinal fluid (temperature ≈4–8°C).The artificial cerebrospinal fluid (ACSF) contained 124 mM NaCl, 26 mM NaHCO3, 2.5 mM KCl, 1.25 mM NaH2PO4, 2.5 mM CaCl2, 1.3 mM MgCl2, and 10 mM glucose. Slices were maintained at room temperature for at least 1 hour in a submerged chamber containing ACSF equilibrated with 95% O2 and 5% CO2 and were then transferred to a super fusion chamber. Field EPSPs were recorded, using microelectrodes (1–3 MΩ) filled with ACSF, at 22° to 25° C. Bipolar stainless steel electrodes were used for the electrical stimulation of Schaffer collaterals and mossy fibers (0.1 ms, 10 to 100 μA pulses, intertrial intervals of 10 to 30 s). Field EPSP recordings of mossy fibers and Schaffer collaterals were taken with a DAM80 amplifier (WPI), under visual control, with an upright microscope (BX50WI, Olympus). Mossy fiber LTP was induced by tetanus at 25 Hz for 5 s, in the presence of 50 μM D-AP5 (D (-) 2-amino-5-phosphonovaleric acid). Mossy fiber responses were identified in wild-type and 152F7 mice with the group 2-metabotropic glutamate receptor selective agonist DCG IV. The inhibitory effects of DCG IV (10 μM) on mossy fiber inputs were similar in mutants and wild-type mice. NBQX (5 μM) was applied at the end of each mossy fiber experiment, to assess the amplitude of fiber volleys. For Schaffer collateral LTP, stimulating and extracellular recording electrodes were placed in the *stratum radiatum*, and the GABA-A receptor antagonist picrotoxin (100 μM) was added to the ACSF. In this series of experiments, the CA1 region was separated from the CA3 region by sectioning the brain slice with a knife before recording. Data were acquired and analyzed blind to genotype for LTP experiments. On- and offline data analyses were carried out with Acquis1-ElPhy software (developed by G. Sadoc, UNIC CNRS and ANVAR, France). Summary data are expressed as means ± SEM. The following drugs were used: NBQX, D-APV, DCGIV (Tocris) and picrotoxin (Sigma).

### Electrophysiological analysis on 189N3 adult mice

All experiments were approved by the Institut National de la Santé et de la Recherche Médicale (INSERM) animal care and use agreement (D-13-055-19) and the European community council directive (2010/63/UE).

#### Preparation of Hippocampal Slices

189N3 KI and WT littermate male mice aged 17–22 week-old were shipped from IGBMC (Strasbourg, France) to INMED (Marseille, France). Mice were deeply anesthetized with xylazine 13 mg/kg / ketamine 66 mg/kg and transcardially perfused with a modified artificial cerebrospinal fluid (mACSF) containing the following (in mM): 132 choline, 2.5 KCl, 1.25 NaH2PO4, 25 NaHCO3, 7 MgCl2, 0.5 CaCl2, and 8 D-glucose prior to decapitation. The brain was then removed rapidly, the hippocampi were dissected, and transverse 450 μM thick slices were cut using a Leica VT1200S vibratome in ice-cold oxygenated (95% O2 and 5% CO2) mACSF. Slices recovered at room temperature for at least 1 h in artificial cerebrospinal fluid (ACSF) containing the following (in mM): 126 NaCl, 3.5 KCl, 1.25 NaH2PO4, 26 NaHCO3, 1.3 MgCl2, 2.0 CaCl2, and 10 D-glucose. Both cutting solution and ACSF were between 290 mOsm and 310 mOsm. All solutions were equilibrated with 95% 2 and 5% CO2, pH 7.4.

#### fEPSP Recordings

Acute slices were individually transferred to a recording chamber maintained at 30–32°C and continuously perfused (2 ml/min) with oxygenated ACSF. Field excitatory postsynaptic potential (fEPSP) were made in the *lucidum* of CA3 area with glass electrodes (2–3 MΩ; filled with normal ACSF) using a DAM-80 amplifier (low filter, 1 Hz; high pass filter, 3 KHz; World Precision Instruments, Sarasota, FL). Mossy fiber-mediated fEPSP were evoked by weak electrical stimulations performed via a bipolar NiCh electrode (NI-0.7F, Phymep, Paris) positioned in the lucidum of CA3 area; the stimulus intensity, pulse duration, and frequency were around 30V, 25μs, and 0.1 Hz, respectively. Data were digitized with a Digidata 1440A (Molecular Devices) to a PC, and acquired using Clampex 10.1 software (PClamp, Molecular Devices). Signals were analyzed off-line using Clampfit 10.1 (Molecular Devices). LTP was induced by tetanus at 25Hz during 5s; D-APV (D-2-amino-5-phosphonovalerate, 40 μM) was included in ACSF during the tetanus to eliminate contamination of MF-CA3 LTP with NMDA receptor-dependent component. At the end of each experiment, 2μM DCG-IV, a group II mGluR agonist, was bath applied to confirm the mossy fiber synaptic origin of fEPSP recorded in CA3. The magnitude of long-term plasticity was determined by comparing baseline-averaged responses before induction with the last 10 min of the experiment. Example traces are averages of at least 30 consecutive sweeps taken from a single representative experiment.

### The Hipposeq analysis

The Hipposeq data, analysis, and website were obtained through collaboration between the Spruston Laboratory (Mark Cembrowski, Nelson Spruston), NeuroSeq (Lihua Wang, Erina Hana, Ken Sugino, Andrew Lemire), and Scientific Computing (Jody Clements, Jonathan Epstein) at the Janelia Research Campus of the Howard Hughes Medical Institute.

#### Statistical analysis

All data are shown as means±SEM. Statistics were performed using IgorPro (Wavemetrics), and statistical significance was determined by the Student’s t test (two tailed distribution, paired) unless otherwise stated.

### Reagents

Stock solutions were prepared in water or DMSO, depending on the manufacturers’ recommendation, and stored at −20°C. Upon experimentation, reagents were bath applied following dilution into ACSF (1/1000). D-APV and DCGIV were purchased from Tocris Bioscience. Salts for making cutting solution and ACSF were purchased from Sigma.

---

**Supplementary Figure 1. Schematic representation of the human YAC 152F7 transgene and of the mouse BAC 189N3 transgene.**

We present here the genomic region of the human YAC 152F7 from the hg38 release (UCSC genome browser).

Note that human YAC 152F7 transgene (570 kb) involves a copy of six hChr21 genes: *RIPPLY3, PIGP, TTC3, DSCR9, VPS26C* and *DYRK1A* respectively.

In contrast, the mouse BAC 189N3 transgene consists of a 152 kb containing only the whole mouse Dyrk1a gene with a 6 kb flanking fragment on the 5’ side and a 19 kb flanking fragment on the 3’ (189N3 BAC clone from Research Genetics).

**Supplementary Figure 2. Schematic representation of the mouse genomic region syntenic of the human YAC 152F7 transgene and that includes the mouse BAC 189N3 transgene.**

We present here the mouse syntenic genomic region of the human YAC 152F7 (Mouse Dec. 2011 (GRCm38/mm10 release) (UCSC genome browser).

The YAC 152F7 transgene (570 kb) involves a copy of six hChr21 genes: *RIPPLY3, PIGP, TTC3, DSCR9, VPS26C* and *DYRK1A* respectively.

In contrast, the syntenic mouse region displays orthologous genes of *RIPPLY3, PIGP, TTC3, DSCR9, VPS26C* and *DYRK1A* in a more compact region of ~300 kb. *DSCR9* is a primate-specific gene (Takamatsu et al., 2002).

**Supplementary Figure 3. Nuclear protein-protein interactions and deregulation of interaction network in transgenic mouse models.**

A. *In situ* proximity ligation assays PLA on primary cortical neurons fixed at DIC7 (red fluorescence) using anti-Ep300 and anti-Smarca2 antibodies, anti-Crebbp and anti-Smarca2 antibodies, anti-Smarca4 and anti-Smarca2 antibodies and anti-Ep300 and anti-Fibrillarin antibodies as a negative control. Nuclear bodies were labelled using Topro3 staining (blue fluorescence). Mean interaction point numbers were calculated in nuclear body of at least 48 cortical neurons at DIC7.

B. *In situ* proximity ligation assays PLA on transgenic 189N3 and WT primary cortical neurons fixed at DIC7 (red fluorescence) using anti-Baf155 and anti-Smarca4 antibodies. Nuclear bodies were labelled using Topro3 staining (blue fluorescence). Mean interaction point numbers were calculated in nuclear body of at least 48 cortical neurons at DIC7.

Scale bars = 10μm. *** p < 0.0005

**Supplementary Figure 4. Quantification of Rims1 RNA in WT and 152F7 hippocampus using quantitative In Situ Hybridation**

False-color image of antisense *Rims1* RNA Q-ISH of hippocampus from juvenile P21 WT and 152F7 mice. Q-ISH was performed using 3H radioactive probes for *Rims1.* Q-ISH quantification indicates a significant down-regulation of *Rims1* in the three subregions of the 152F7 mouse hippocampus

**Supplementary Figure 5. Comparison of transcripts encoding presynaptic proteins using either Q-RT PCR from laser assisted microdissection or RNAseq single cell transcriptomics**

A. Regions of interest from which RNAseq single cell transcriptomics was performed in population(s) of excitatory cells is indicated (The Hipposeq data obtained from Janelia Farm website) (Cembrowski et al., 2016)

B. Comparison of our data from Laser-assisted microdissection of the three subregions of P21 mouse hippocampus of juvenile transgenic 152F7 mice compared to their WT siblings (see Figure 3D) and RNAseq data for *Rims1, Syn2, Rab3a* and *Munc13-1* transcripts.

Scale bar = 1mm. *p<0.01**p<0.001. ***p<0.0001

