## Supplementary Figures for "DYRK1A up-regulation specifically impairs a presynaptic form of long-term potentiation"

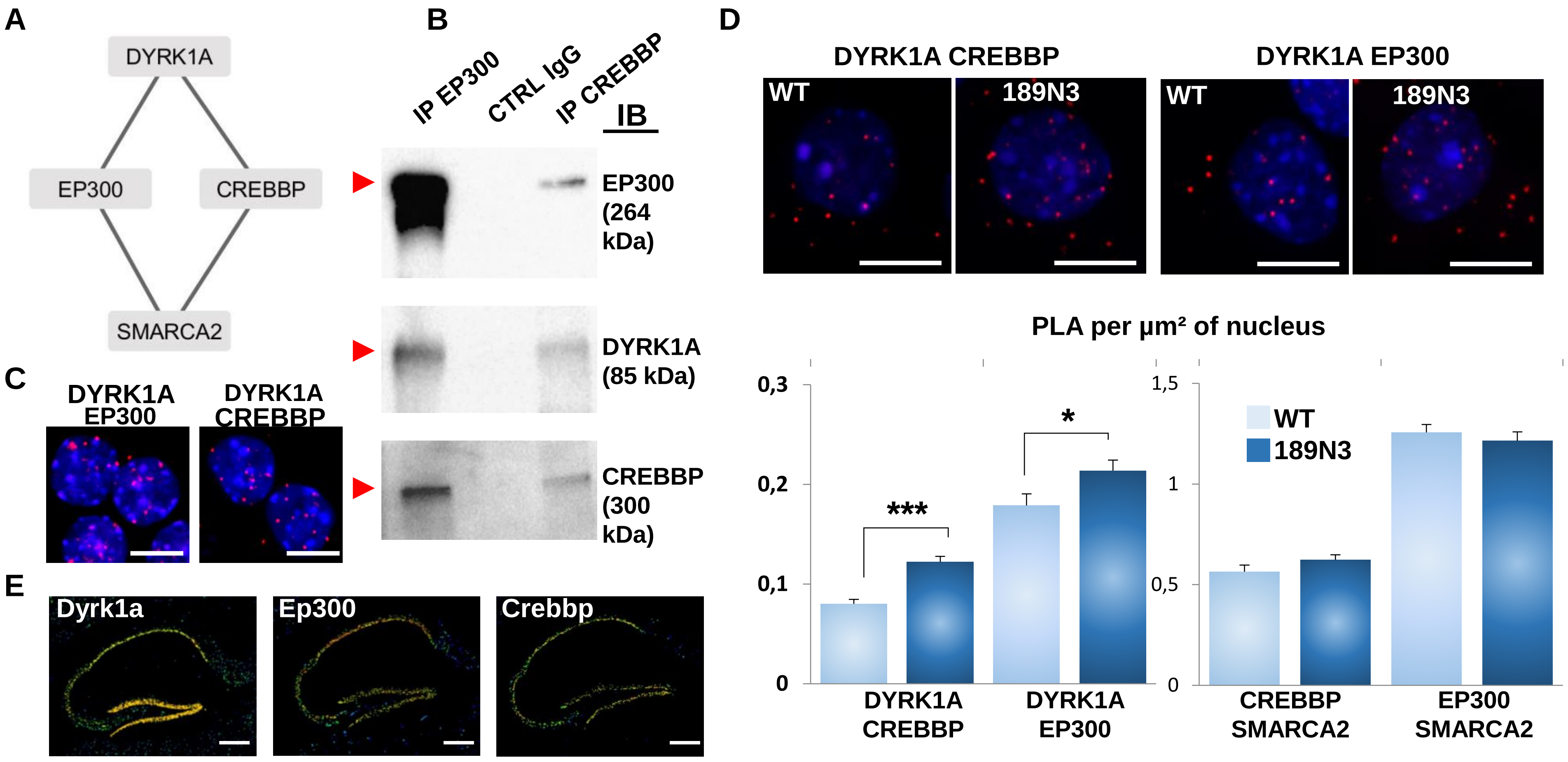
A
DYRK1A
EP300
CREBBP
SMARCA2
B
IP EP300
CTRL IgG
IP CREBBP
IB
EP300 (264 kDa)
DYRK1A (85 kDa)
CREBBP (300 kDa)
C
DYRK1A EP300
DYRK1A CREBBP
D
DYRK1A CREBBP
DYRK1A EP300
WT
189N3
WT
189N3
PLA per µm² of nucleus
0,3
0,2
0,1
0
1,5
1
0,5
0
***
*
WT
189N3
DYRK1A CREBBP
DYRK1A EP300
CREBBP SMARCA2
EP300 SMARCA2
E
Dyrk1a
Ep300
Crebbp

Figure 1

**Figure 1. Interaction of HSA21 DYRK1A with chromatin remodelers**

A. Schematic representation of DYRK1A interaction with EP300 and CREBBP. B. HEK293 cells were immunoprecipitated (IP) using anti-EP300 and anti-CREBBP antibodies and using anti-IgG antiboby as a negative control. The input and precipitated fractions were analyzed by western blot using anti-Ep300, anti-Dyrk1a and anti-Crebbp antibodies. The arrows indicate proteic bands at the expected size. Note that no cross-reaction was found with the IgGs.

C-D. *In situ* proximity ligation assays (PLA) on primary cortical neurons fixed at DIC7 (red fluorescence) using anti-Dyrk1a and anti-Ep300 or anti-Crebbp, anti-Smarca2 and anti-Ep300 or anti-Crebbp antibodies. Nuclear bodies were labelled using Topro3 staining (blue fluorescence). Mean interaction point numbers were calculated in nuclear body of 45 to 89 cortical neurons at DIC7 (from 3 to 5 different embryos per genotype). PLA using anti-Ep300 and anti-Fibrillarin antibodies were performed as a negative control and no difference was shown between transgenic 189N3 and WT cortical neurons. Scale bars=10µm. * $p < 0.05$; *** $p < 0.0005$. E. False-color image of ISH from Allen Brain Atlas showing Dyrk1a, Ep300 and Crebbp transcript expression in adult mouse hippocampus. Scale bar = 100µm.

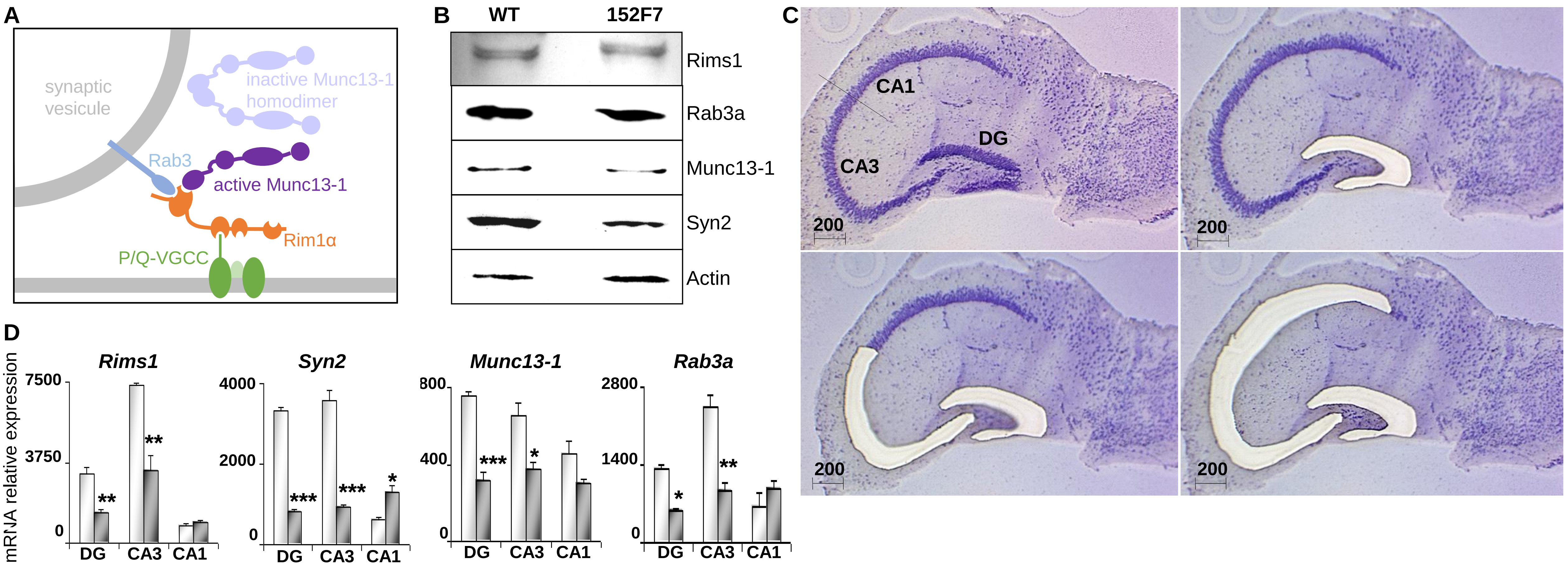
A
synaptic
vesicule
inactive Munc13-1
homodimer
Rab3
active Munc13-1
Rim1α
P/Q-VGCC
B
WT
152F7
Rims1
Rab3a
Munc13-1
Syn2
Actin
C
CA1
CA3
DG
200
200
200
200
D
mRNA relative expression
Rims1
7500
3750
0
**
**
DG
CA3
CA1
Syn2
4000
2000
0
***
***
*
DG
CA3
CA1
Munc13-1
800
400
0
***
*
DG
CA3
CA1
Rab3a
2800
1400
0
*
**
DG
CA3
CA1

Figure 2

**Figure 2. Presynaptic protein expression in the adult 152F7 mouse hippocampus as compared to control mice.**

A. Schematic representation of molecules involved in glutamate release from presynaptic vesicles.
B. Western blotting of Rim1, Rab3a, Munc13-1 and Synapsin 2A proteins from hippocampus extracts of wild-type and 152F7 transgenic mice. The amounts of protein were normalised to actin levels. Four to five hippocampi were pooled. There was a decrease in the amount of Synapsin 2A in the 152F7 transgenic mice (152F7) as compared to the wild type (WT).
C & D. Laser-assisted microdissection of the three subregions of P21 mouse hippocampus stained with toluidine blue (C). Scale bar = 200 µm. From Laser-assisted microdissection of the three subregions of P21 mouse hippocampus, *Rims1, Syn2, Rab3a* and *Munc13-1* transcripts are down regulated in the DG and CA3 hippocampal subregions of juvenile transgenic 152F7 mice compared to their WT siblings as shown by Q-RT-PCR analysis (D).
Scale bar = 1mm. *$p$<0.01**$p$<0.001. ***$p$<0.0001

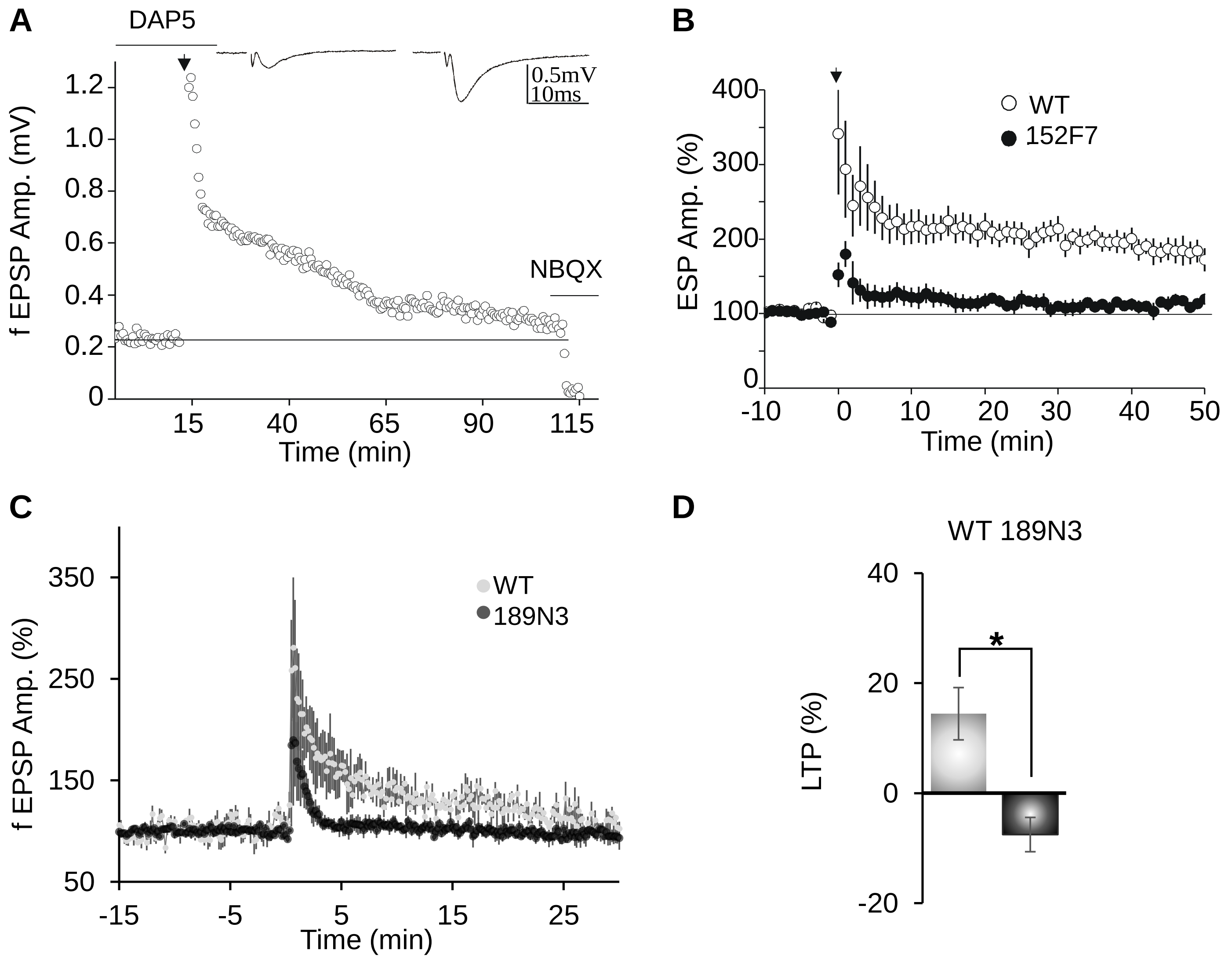
A
DAP5
NBQX
0.5mV
10ms
f EPSP Amp. (mV)
1.2
1.0
0.8
0.6
0.4
0.2
0
15
40
65
90
115
Time (min)
B
WT
152F7
ESP Amp. (%)
400
300
200
100
0
-10
0
10
20
30
40
50
Time (min)
C
WT
189N3
f EPSP Amp. (%)
350
250
150
50
-15
-5
5
15
25
Time (min)
D
WT 189N3
LTP (%)
40
20
0
-20
*

Figure 3

**Figure 3. Presynaptic LTP between Dentate Gyrus mossy fibers and CA3 impaired in both adult 152F7 and 189N3 mouse hippocampus as compared to control mice.**

A. Example of a temporal trace of mossy fiber LTP in a wild type mouse. Mossy fiber LTP was induced by a single tetanus of 25 Hz (for 5 s, black arrow) in the presence of 50 µM of DAP 5. NBQX was applied at the end to obtain information concerning the fiber volley. Inset: example traces of a baseline fEPSP (left) and fEPSP after LTP (right).
B. Summary of mossy fiber LTP in transgenic 152F7 compared to WT mice, showing LTP induction defect in the DS mouse model. Data are means+/ SEM and calculated from 4 different mice with 3 slices per mouse for both genotypes.
C. D. Summary of mossy fiber LTP in transgenic 189N3 compared to WT mice (**C**). Average fEPSP amplitude 20 min after LTP induction in WT versus 189N3 mice, showing LTP induction defect in the DS mouse model (D). Recording were performed on 4 to 6 mice in each genotypes. Data are means+/ SEM.

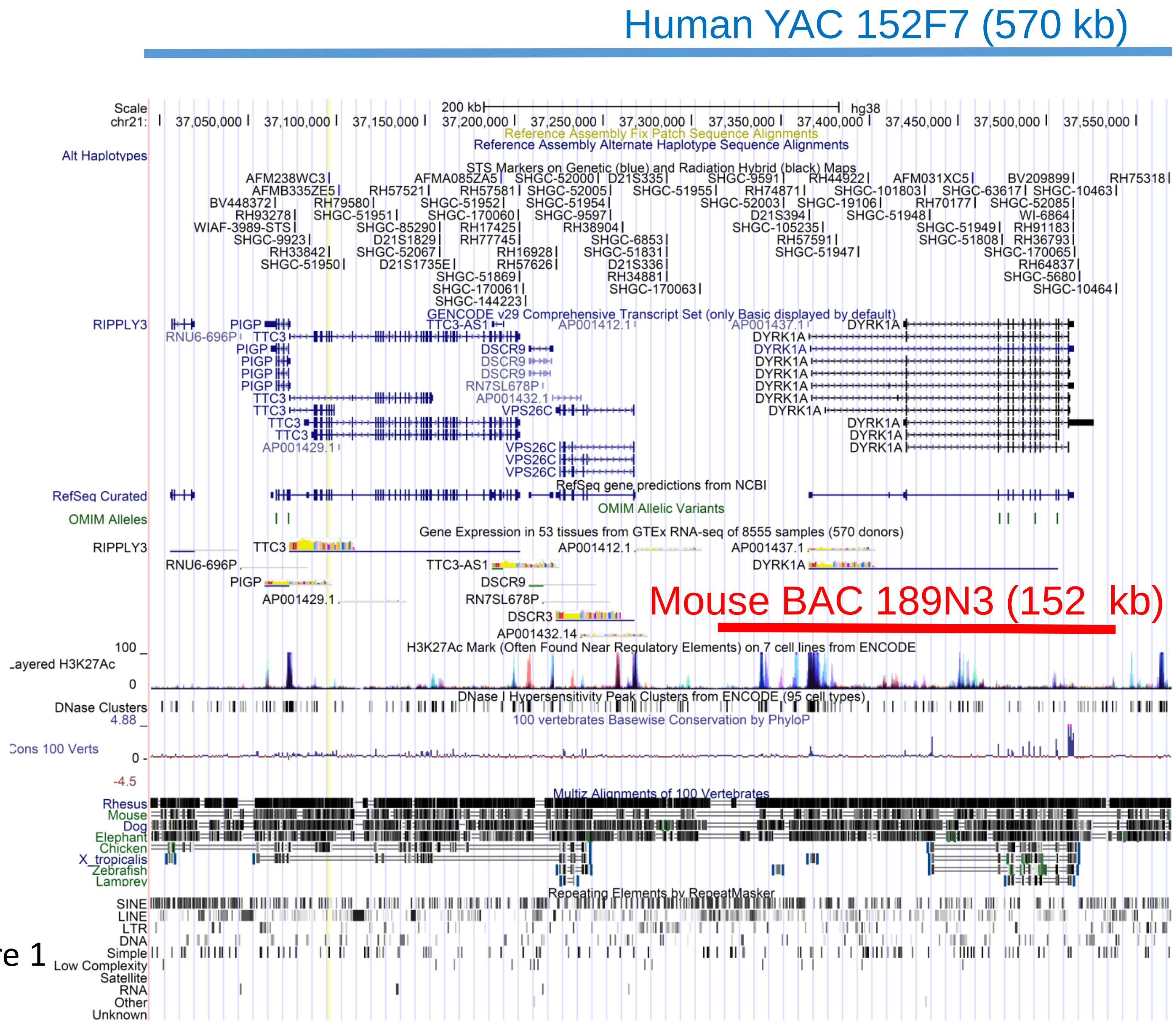


Supplementary Figure 1

**Supplementary Figure 1. Schematic representation of the human YAC 152F7 transgene and of the mouse BAC 189N3 transgene.**

We present here the genomic region of the human YAC 152F7 from the hg38 release (UCSC genome browser).
Note that human YAC 152F7 transgene (570 kb) involves a copy of six hChr21 genes: *RIPPLY3, PIGP, TTC3, DSCR9, VPS26C* and *DYRK1A* respectively.
In contrast, the mouse BAC 189N3 transgene consists of a 152 kb containing only the whole mouse Dyrk1a gene with a 6 kb flanking fragment on the 5' side and a 19 kb flanking fragment on the 3‘ (189N3 BAC clone from Research Genetics).

Supplementary Figure 2

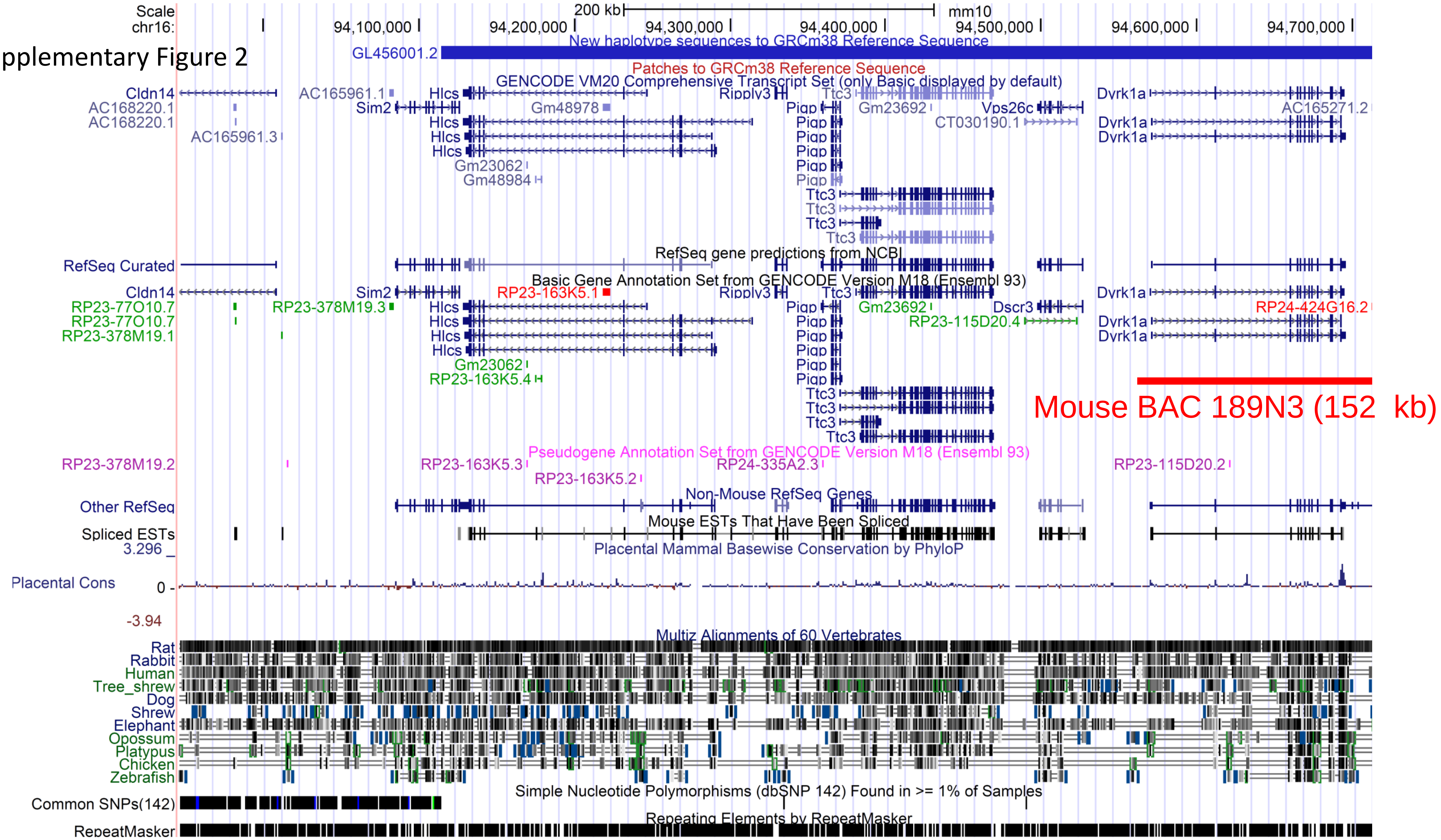

**Supplementary Figure 2. Schematic representation of the mouse genomic region syntenic of the human YAC 152F7 transgene and that includes the mouse BAC 189N3 transgene.**

We present here the mouse syntenic genomic region of the human YAC 152F7 (Mouse Dec. 2011 (GRCm38/mm10 release) (UCSC genome browser).

The YAC 152F7 transgene (570 kb) involves a copy of six hChr21 genes: *RIPPLY3, PIGP, TTC3, DSCR9, VPS26C* and *DYRK1A* respectively.

In contrast, the syntenic mouse region displays orthologous genes of *RIPPLY3, PIGP, TTC3, DSCR9, VPS26C* and *DYRK1A* in a more compact region of ~300 kb. *DSCR9* is a primate-specific gene (Takamatsu et al., 2002).

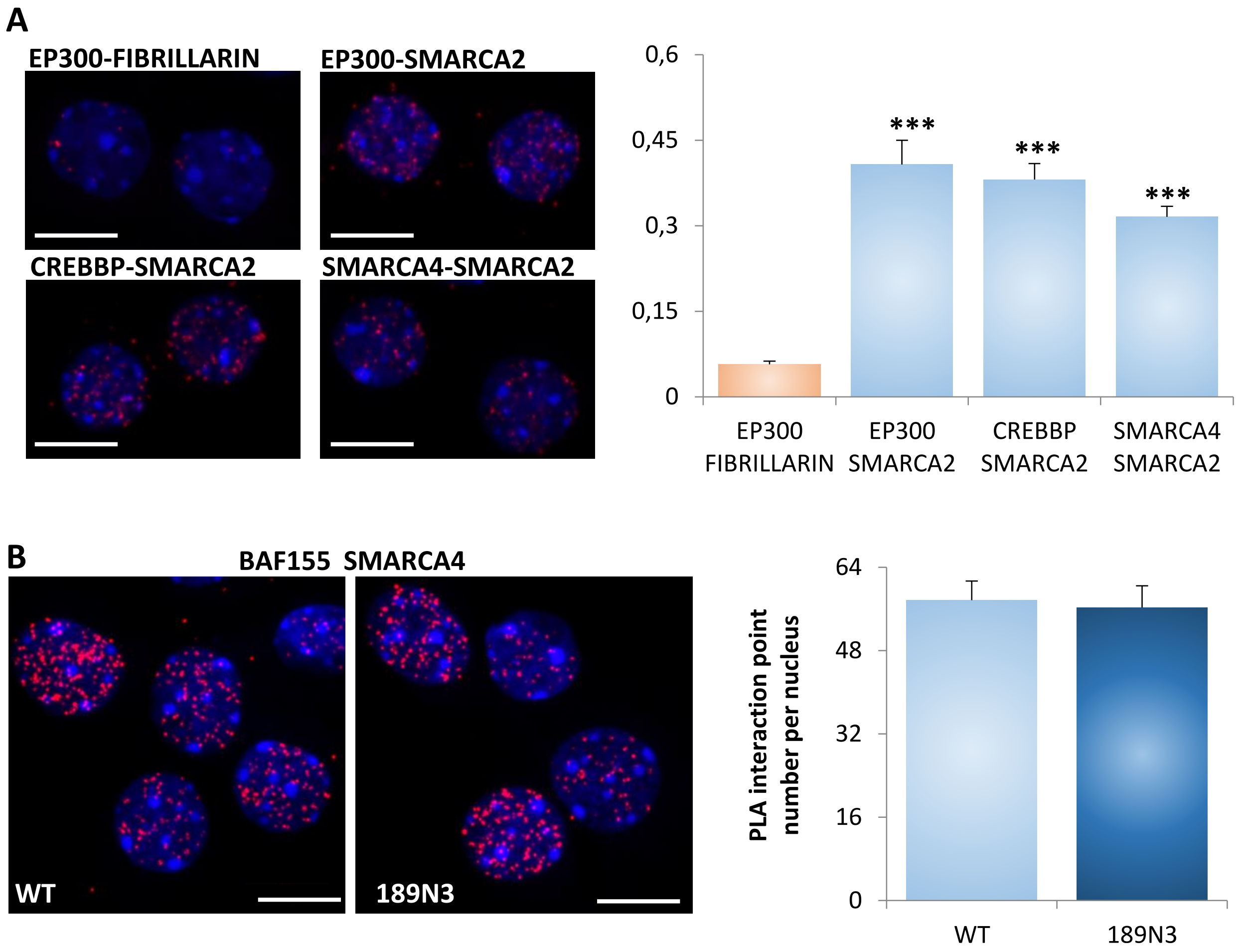


Supplementary Figure 3

**Supplementary Figure 3. Nuclear protein-protein interactions and deregulation of interaction network in transgenic mouse models.**

A. *In situ* proximity ligation assays PLA on primary cortical neurons fixed at DIC7 (red fluorescence) using anti-Ep300 and anti-Smarca2 antibodies, anti-Crebbp and anti-Smarca2 antibodies, anti-Smarca4 and anti-Smarca2 antibodies and anti-Ep300 and anti-Fibrillarin antibodies as a negative control. Nuclear bodies were labelled using Topro3 staining (blue fluorescence). Mean interaction point numbers were calculated in nuclear body of at least 48 cortical neurons at DIC7.
B. *In situ* proximity ligation assays PLA on transgenic 189N3 and WT primary cortical neurons fixed at DIC7 (red fluorescence) using anti-Baf155 and anti-Smarca4 antibodies. Nuclear bodies were labelled using Topro3 staining (blue fluorescence). Mean interaction point numbers were calculated in nuclear body of at least 48 cortical neurons at DIC7.
Scale bars = 10μm. *** $p < 0.0005$

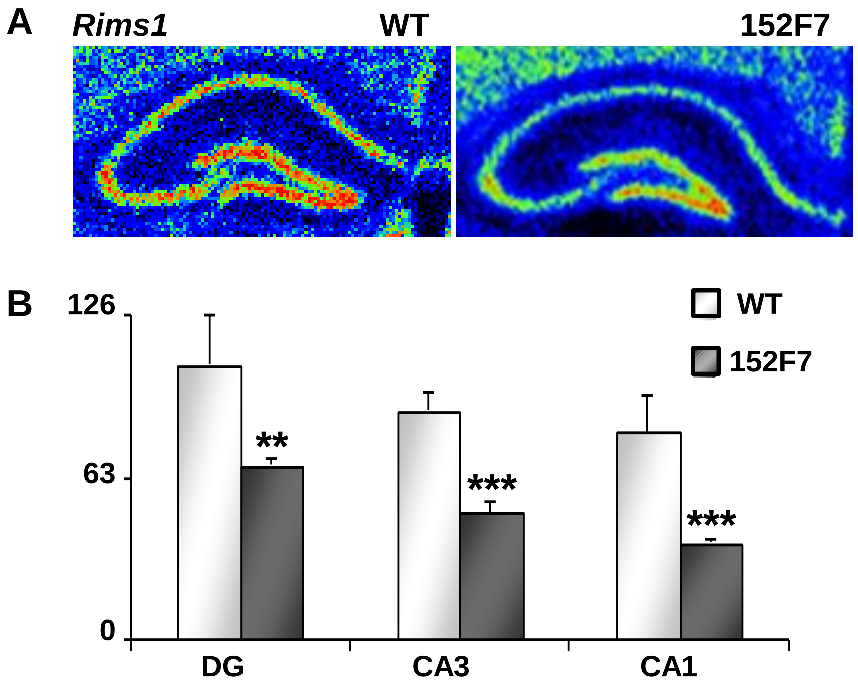


Supplementary Figure 4

**Supplementary Figure 4. Quantification of Rims1 RNA in WT and 152F7 hippocampus using quantitative In Situ Hybridation**

False-color image of antisense *Rim*s1 RNA Q-ISH of hippocampus from juvenile P21 WT and 152F7 mice. Q-ISH was performed using 3H radioactive probes for *Rims1*. Q-ISH quantification indicates a significant down-regulation of *Rims1* in the three subregions of the 152F7 mouse hippocampus.

A

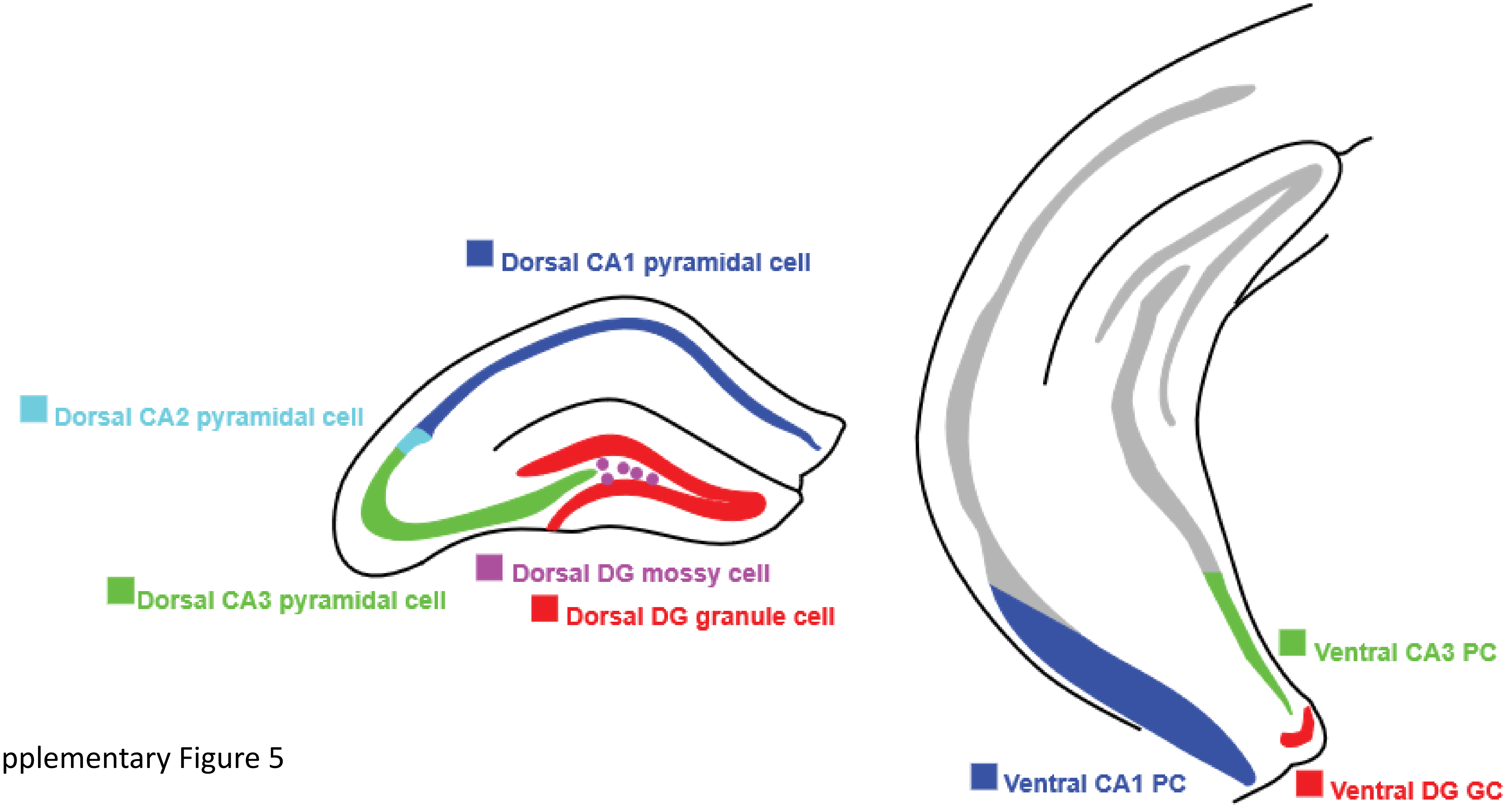


Supplementary Figure 5

**Supplementary Figure 5. Comparison of transcripts encoding presynaptic proteins using either Q-RT PCR from laser assisted microdissection or RNAseq single cell transcriptomics**

A. Regions of interest from which RNAseq single cell transcriptomics was performed in population(s) of excitatory cells is indicated (The Hipposeq data obtained from Janelia Farm website) (Cembrowski et al., 2016)
B. Comparison of our data from Laser-assisted microdissection of the three subregions of P21 mouse hippocampus of juvenile transgenic 152F7 mice compared to their WT siblings (see Figure 3D) and RNAseq data for *Rims1, Syn2, Rab3a* and *Munc13-1* transcripts.
Scale bar = 1mm. *$p$<0.01**$p$<0.001. ***$p$<0.0001

Supplementary Figure 6

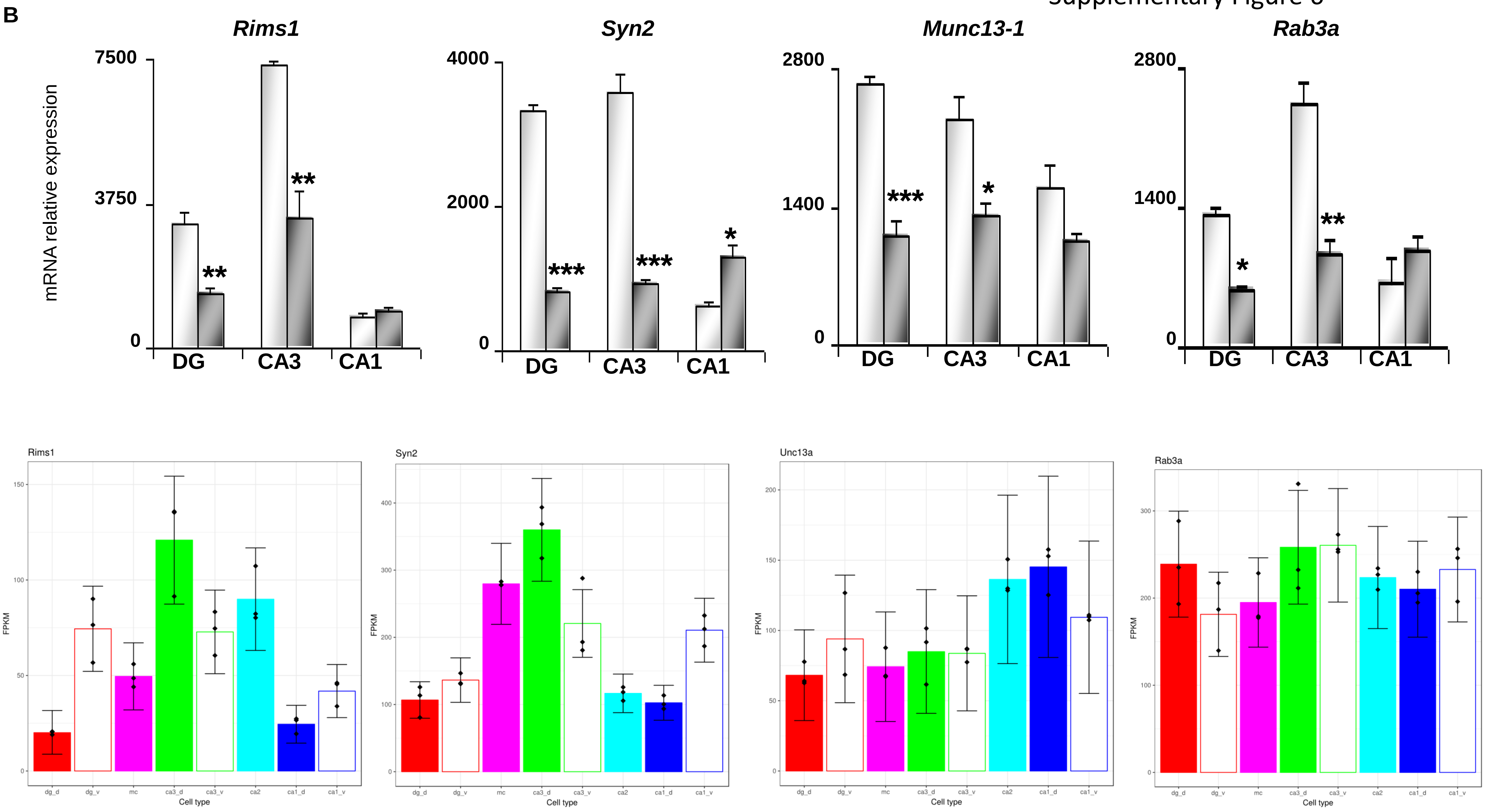

**Supplementary Figure 5. Comparison of transcripts encoding presynaptic proteins using either Q-RT PCR from laser assisted microdissection or RNAseq single cell transcriptomics**

A. Regions of interest from which RNAseq single cell transcriptomics was performed in population(s) of excitatory cells is indicated (The Hipposeq data obtained from Janelia Farm website) (Cembrowski et al., 2016)
B. Comparison of our data from Laser-assisted microdissection of the three subregions of P21 mouse hippocampus of juvenile transgenic 152F7 mice compared to their WT siblings (see Figure 3D) and RNAseq data for *Rims1, Syn2, Rab3a* and *Munc13-1* transcripts.
Scale bar = 1mm. *$p$<0.01**$p$<0.001. ***$p$<0.0001
